# PandaDock: An Open-Source Molecular Docking Platform with Flexible-Ligand Search and Equivariant Neural Scoring

**DOI:** 10.64898/2026.08.19.745667

**Authors:** Pritam Kumar Panda

**Affiliations:** Department of Anesthesiology, Perioperative and Pain Medicine, Stanford University, Stanford, CA 94305, USA

**Keywords:** molecular docking, flexible-ligand docking, scoring functions, graph neural networks, SE(3)-equivariance, open-source software

## Abstract

We present PandaDock, an open-source molecular docking platform implementing flexible-ligand conformational search with analytic gradients, a precomputed affinity grid engine, specialized modules for induced-fit, metal-coordination and tethered docking, and an SE(3)-equivariant graph neural network scoring function trained at scale. Ligand flexibility is represented as a torsion tree and pose parameters are optimized by Monte Carlo with Metropolis acceptance refined by L-BFGS, with rotational gradients obtained in closed form through the derivative of the SO(3) exponential map rather than by finite differences. Affinity grids are built by a blocked neighbor-selection scheme that is exact and 5.6–9.7*×* faster than dense evaluation, and may be cached across ligands sharing a receptor and site, reducing a six-ligand series from 29.3 s to 10.4 s. On 814 protein–ligand complexes spanning 14 target families, PandaDock recovers a pose within 2 Å of the crystal geometry in 33.7% of cases at rank 1 and in 57.0% of cases within the returned ensemble. The GNN scoring function is trained on 741,706 co-folded complexes from SAIR under target-disjoint splits, reaching a Pearson *r* of 0.407 on 90,219 held-out complexes and transferring to 202 independent crystal structures with measured *K*_*i*_, *K*_*d*_, IC_50_ or EC_50_ at *r* = 0.467. We report the model against three controls, a target-mean predictor, a ligand-descriptor-only baseline, and within-target correlations, and document both where it performs and where it does not, including its unsuitability for pose rescoring. On an independent 30-compound series against a single GABA_*A*_ receptor target, PandaDock’s empirical scoring function ranks 8th of 25 methods evaluated, ahead of every AutoDock Vina and Vinardo configuration tested, while the GNN scores below Vina, consistent with the within-target ceiling identified on SAIR. At full scale on the PDBbind v2020 refined set (n = 4,640, native crystal poses), the fully independent SAIR model reaches *r* = 0.531, and a dedicated model trained on PDBbind alone under a target-disjoint split reaches *r* = 0.690 on its own held-out test complexes – the strongest evidence in this work that PandaDock’s affinity predictions generalize. PandaDock is distributed under an open-source license at https://github.com/pritampanda15/PandaDock with a complete command-line interface and a reproducible benchmarking harness.

## 1 Introduction

Molecular docking is a cornerstone of structure-based drug discovery, enabling virtual screening of large compound libraries and structure-guided lead optimization [Kitchen et al., 2004; Meng et al., 2011]. A docking calculation comprises two components: a conformational search generating plausible binding poses, and a scoring function ranking them and estimating binding affinity [Sousa et al., 2006], both remain active areas of research.

Established open-source platforms i.e., AutoDock Vina [Trott and Olson, 2010], smina [Koes et al., 2013] and GNINA [McNutt et al., 2021] have made docking broadly accessible, while GPU-accelerated implementations [Santos-Martins et al., 2021; Tang et al., 2022] have extended it to billion-compound libraries. Learned scoring functions, from convolutional networks on voxelised binding sites [Ragoza et al., 2017] to graph neural networks operating directly on molecular graphs [Lim et al., 2019; Jiang et al., 2021], offer improved affinity correlation, and equivariant architectures [Thomas et al., 2018; Satorras et al., 2021] enforced by constructing the rotational and translational invariance that binding energy must physically obey. PandaDock is an open-source platform combining a from-scratch flexible-ligand search engine with a large-scale learned scoring function, designed so that each component can be evaluated independently. Our contributions are: (i) **Flexible-ligand search with analytic gradients**. A torsion-tree representation optimized by Monte Carlo with L-BFGS refinement, in which rotational gradients are computed in closed form through the SO(3) exponential map rather than by finite differences. (ii) **An exact fast grid engine**. A blocked neighbor-selection scheme for affinity grid construction that is provably lossless and 5.6–9.7*×* faster, with a signature-keyed cache that amortizes grid cost across a screening campaign. (iii) **Learned scoring at scale**. An SE(3)-equivariant heterogeneous GNN trained on 741,706 co-folded complexes under target-disjoint splits, validated on independent experimental crystal structures, on a single-target prospective series (Section 3.5), and at full scale on the PDBbind v2020 refined set (Section 3.6). (iv) **Specialized docking modules**. Induced-fit, metal-coordination and tethered docking, exposed through a unified command-line interface.(v) **A reproducible benchmarking harness** covering 814 complexes across 14 target families, with evaluation controls that separate pose selection from conformational sampling and ligand ranking from target ranking.

## 2 Methods

### 2.1 Platform architecture

PandaDock is organized into four layers. A *preprocessing* layer prepares receptors and ligands, splitting complexes using the PDB Chemical Component Dictionary to identify the biologically relevant ligand and to distinguish it from crystallization additives and polymer residues. A *search* layer implements the torsion tree, the objective function and its gradients, and the affinity grid engine. A *scoring* layer provides the empirical Vina-form function and the GNN. An *analysis* layer provides symmetry-corrected RMSD, interaction fingerprinting and report generation. Each layer is usable independently through the Python API or the command-line interface (Section 2.9). Figure 1 summarizes the workflow and the scoring network.

**Figure 1:**
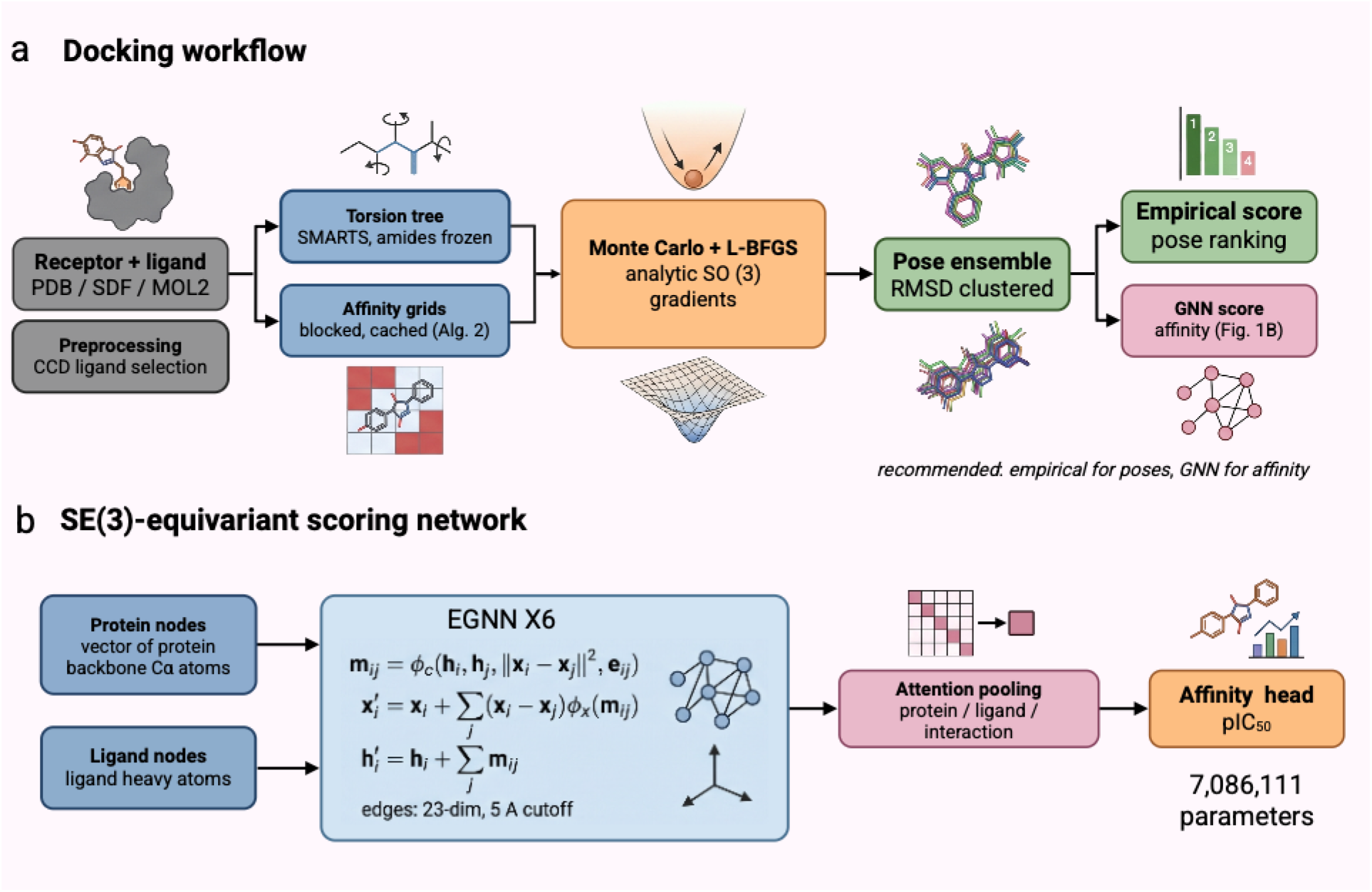
PandaDock architecture. (a) Docking workflow: receptor and ligand are preprocessed, the ligand is converted to a torsion tree and the receptor to blocked affinity grids, poses are optimized by Monte Carlo with L-BFGS refinement using analytic SO(3) gradients, and the resulting ensemble is clustered by symmetry-corrected RMSD. Pose ranking uses the empirical function; the GNN estimates affinity for a selected pose. (b) SE(3)-equivariant scoring network: protein and ligand atoms enter as separate node types, six EGNN layers perform equivariant message passing, and attention pooling feeds an affinity regression head.

### 2.2 Ligand representation

#### 2.2.1 Torsion tree

Ligand flexibility is represented as a torsion tree. Rotatable bonds are identified with the SMARTS pattern

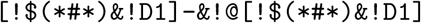

which excludes terminal atoms, triple bonds and ring bonds. Amide C–N bonds are additionally frozen, since their partial double-bond character makes free rotation physically incorrect. The tree is rooted on the largest rigid fragment, minimizing the number of atoms displaced by any single torsion rotation.

A pose is parameterized by

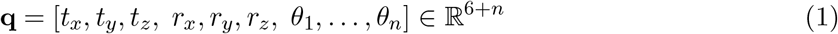

where **t** is a translation, **r** a rotation in axis–angle (exponential-map) form and *θ*_*i*_ the torsion angles. Torsional degrees of freedom are capped at 32, retaining those closest to the root; ligands exceeding the cap are reported rather than silently truncated. Coordinates are generated by traversing the tree from the root, applying each torsion rotation to its sub-tree, then applying the global rotation and translation:

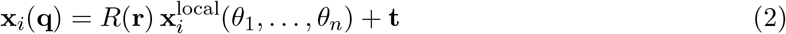

#### 2.2.2 Analytic rotational gradients

Local optimization requires *∂E/∂***q**. For the rotational parameters, finite differences require six additional energy evaluations per step and introduce step-size sensitivity. We instead differentiate the SO(3) exponential map in closed form [Gallego and Yezzi, 2015]. For a rotation vector **r** with *θ* = ∥**r**∥, the derivative of the rotation matrix with respect to component *r*_*k*_ is

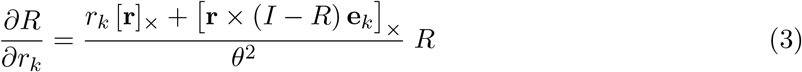

where [·]_×_ denotes the skew-symmetric cross-product matrix and **e**_*k*_ the *k*th basis vector. The small-angle limit is handled separately to avoid the removable singularity as *θ* → 0. Torsional gradients are obtained by projecting per-atom forces onto the rotation axis of each rotatable bond. Uniform random rotations for search initialization are drawn by Shoemake’s method [Shoemake, 1992], with the quaternion negated when *w <* 0 so that the recovered rotation angle lies in [0, *π*].

### 2.3 Conformational search

Search proceeds by independent Monte Carlo runs with Metropolis acceptance, each followed by L-BFGS local refinement (Algorithm 1). Exhaustiveness, the number of independent runs defaults to:

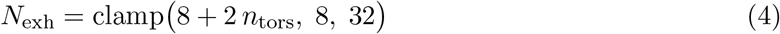

so that flexible ligands receive sampling proportional to their degrees of freedom rather than a fixed budget. Returned poses are clustered by symmetry-corrected RMSD at 2 Å, so that the output represents distinct binding modes rather than near-duplicates.

#### Algorithm 1

Flexible-ligand docking

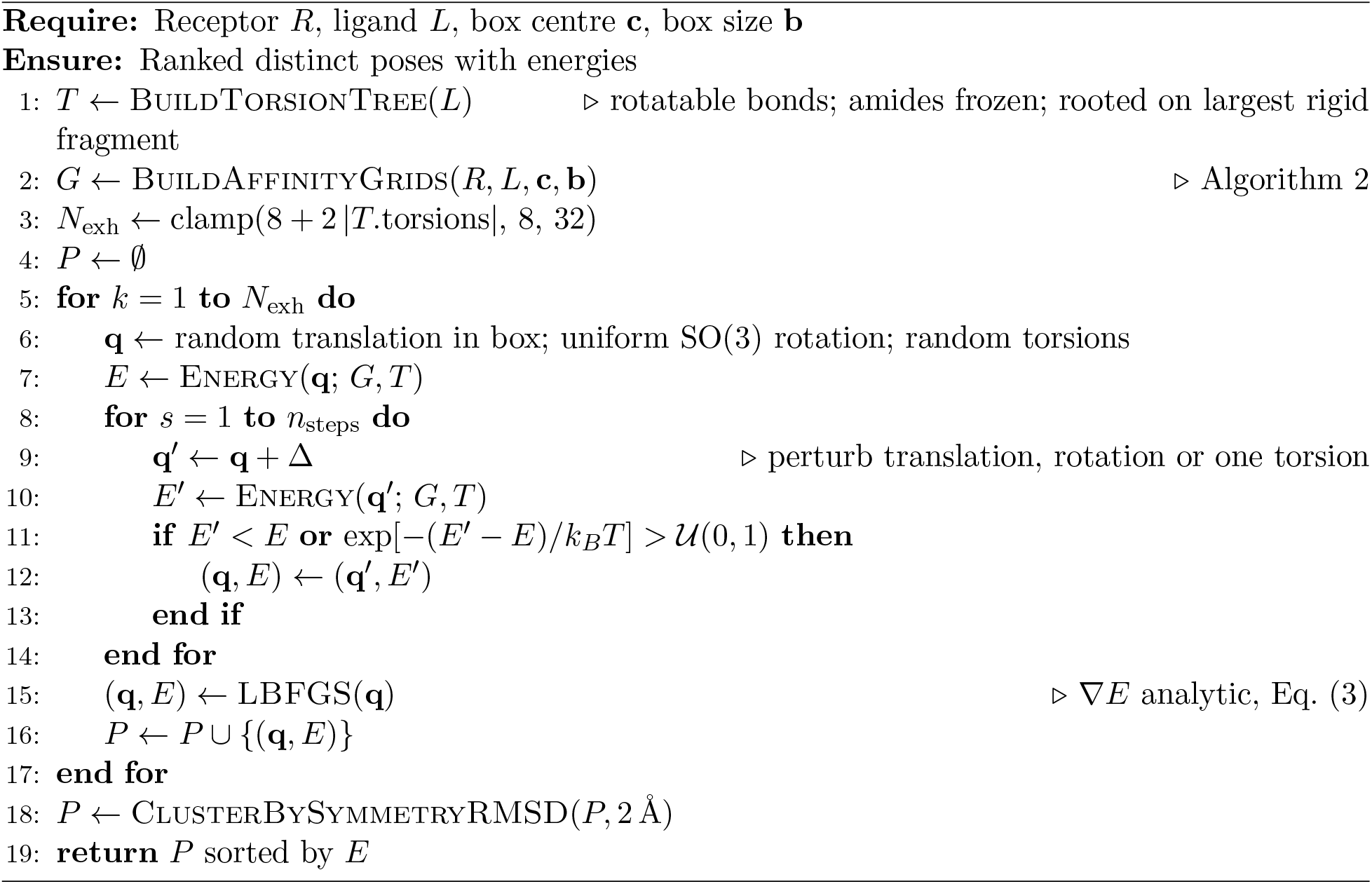

### 2.4 Scoring function and affinity grids

#### 2.4.1 Empirical scoring

The empirical scoring function follows the AutoDock Vina functional form [Trott and Olson, 2010], evaluated on the surface distance 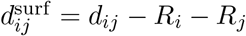:

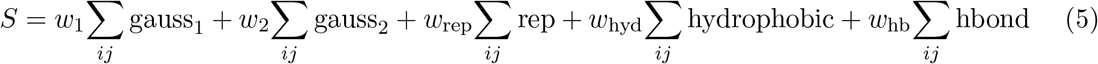

with a torsional penalty applied to the final ranked energy. Receptor atom typing is residue-aware: a flat element-to-type mapping would type backbone carbonyl carbons as `C.3` rather than `C.2` and aromatic ring carbons as `C.3` rather than `C.ar`, which materially changes the features subsequently presented to the learned scoring function.

#### 2.4.2 Blocked grid construction

Receptor contributions are precomputed on grids at 0.375 Å spacing – one grid per distinct ligand atom *signature*, the tuple (radius, hydrophobic, donor, acceptor) – and interpolated trilinearly during search. Grid construction dominates the cost of short runs: profiling attributed 51% of a representative docking run to it.

A dense implementation evaluates every energy term for every grid point against every receptor atom and applies the distance cutoff last. Since an atom contributes only where

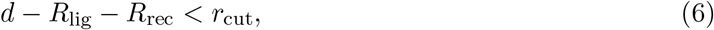

atoms further than 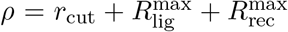 from a region of space contribute exactly zero there. Algorithm 2 exploits this by walking the grid in compact three-dimensional blocks and selecting, per block, only the atoms within reach. The exclusion is exact rather than a heuristic truncation, so the resulting grids are bit-identical to dense evaluation.

##### Algorithm 2

Blocked affinity grid construction with signature cache

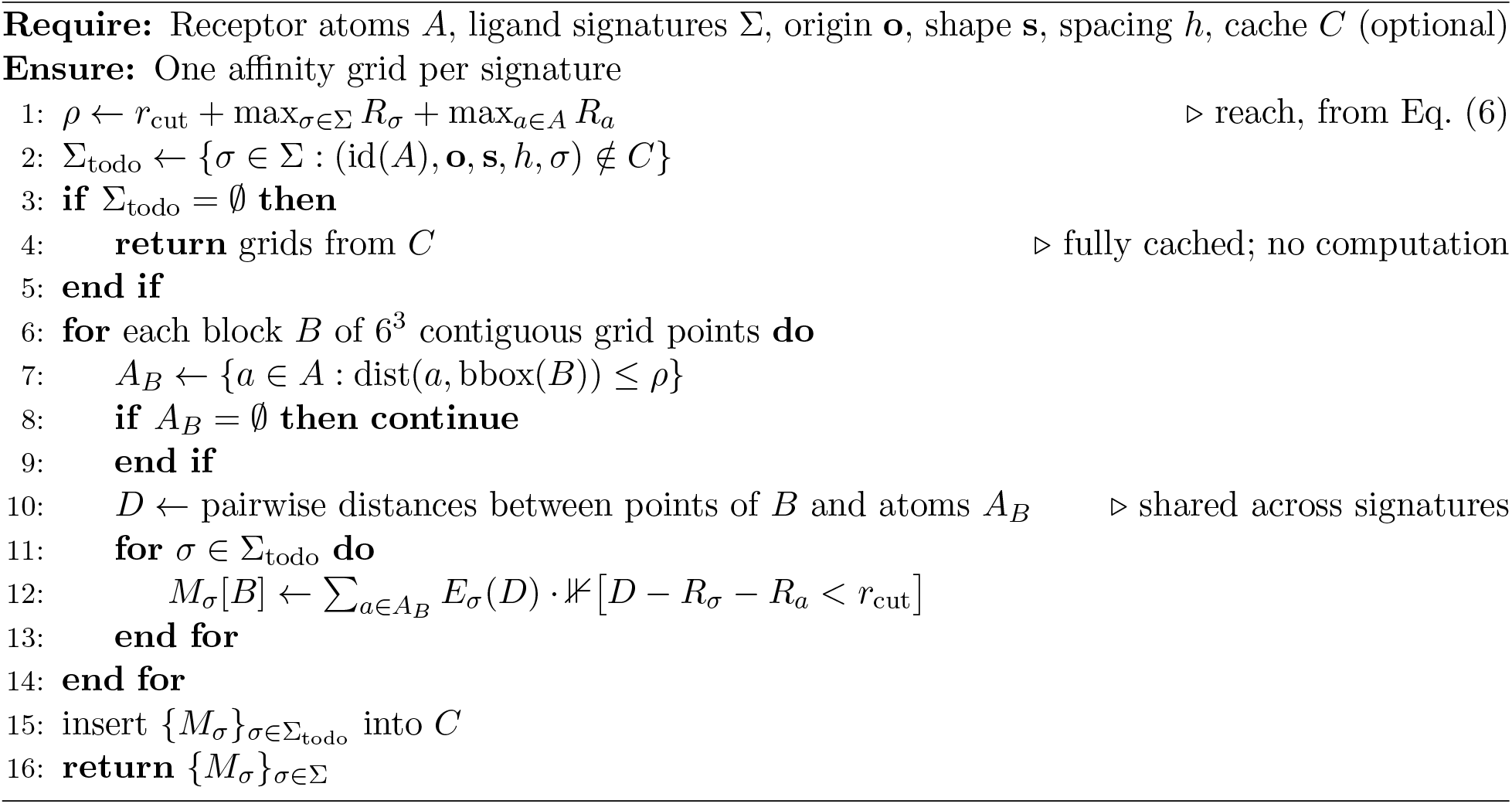

#### 2.4.3 Grid caching for virtual screening

A grid depends on the receptor, the box and an atom signature, but not on ligand identity. Grids may therefore be reused across every ligand docked into the same site such as the separation that `autogrid` provides for AutoDock, and whose absence causes a screening campaign to pay for identical grids once per ligand. PandaDock exposes a signature-keyed cache (Algorithm 2, lines 2–5). The key covers receptor coordinates and radii, grid origin, shape and spacing, so a different receptor, a translated receptor or a different box all correctly miss, and a ligand introducing new atom types builds only the grids it adds.

### 2.5 SE(3)-equivariant GNN scoring function

#### 2.5.1 Graph representation

Complexes are represented as heterogeneous graphs *G* = (*V, E*) with protein and ligand node types. Protein nodes are heavy atoms within 10 Å of the ligand centroid; ligand nodes are all ligand heavy atoms. Node features (Table 1) are 56-dimensional and edge features (Table 2) 23-dimensional. Four edge types are defined: protein→ligand and ligand→protein interactions within 5 Å, and intramolecular contacts within each molecule.

**Table 1:** Node feature composition (56 dimensions).

| Feature | Dims | Description |
| --- | --- | --- |
| Element type | 10 | One-hot (C, N, O, S, P, H, F, Cl, Br, Other) |
| SYBYL atom type | 16 | Compressed embedding of $\sim 26$ types |
| Hybridization | 4 | sp, sp<sup>2</sup>, sp<sup>3</sup>, other |
| H-bond donor/acceptor | 2 | Binary flags |
| Aromaticity, ring | 2 | Binary flags |
| Partial charge | 1 | Clipped to $[-1, 1]$ |
| Residue type | 21 | One-hot; zero for ligand atoms |

**Table 2:**
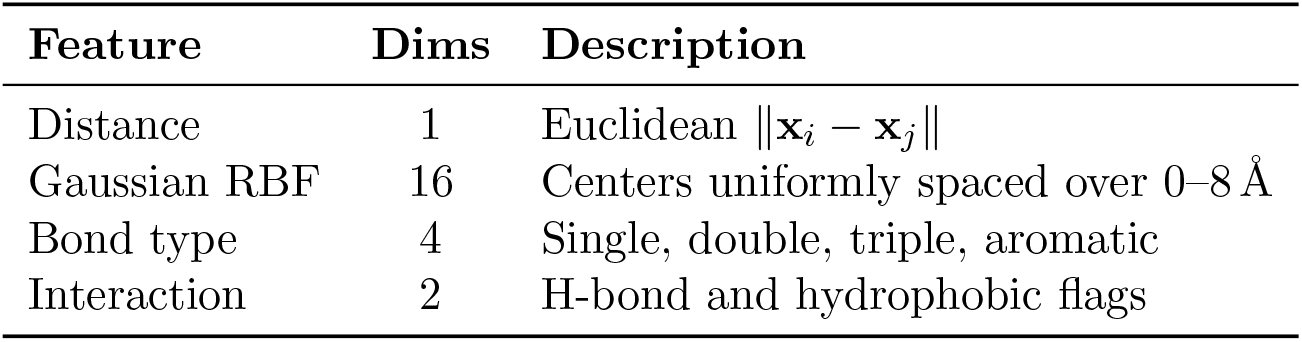
Edge feature composition (23 dimensions).

| Feature | Dims | Description |
| --- | --- | --- |
| Distance | 1 | Euclidean $\ \mathbf{x}_i - \mathbf{x}_j\ $ |
| Gaussian RBF | 16 | Centers uniformly spaced over 0–8 Å |
| Bond type | 4 | Single, double, triple, aromatic |
| Interaction | 2 | H-bond and hydrophobic flags |

Feature construction is the dominant per-sample cost during training. Every feature except partial charge is a pure function of six discrete attributes, and a full dataset contains only 364 distinct combinations, so the discrete portion is memoised exactly. Combined with replacing a scalar `np.clip` by branches, this raises throughput from 446 to 787 samples/s per worker with bit-identical output.

#### 2.5.2 Network architecture

The network follows the E(*n*)-equivariant GNN framework [Satorras et al., 2021]. A physically meaningful scoring function must satisfy

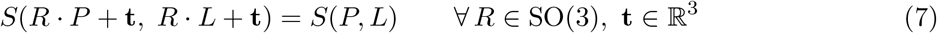

which the architecture enforces by construction. Each layer computes

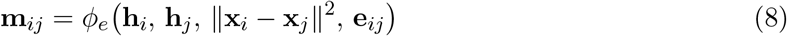

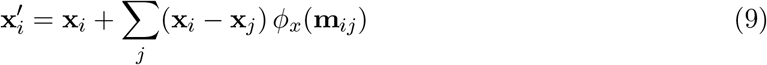

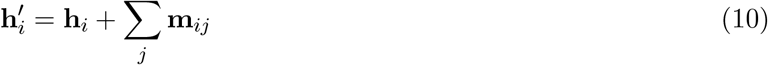

Six such layers at hidden dimension 256 are followed by attention pooling over protein and ligand nodes, concatenation with their element-wise interaction, and an affinity regression head. The model has 7,086,111 parameters.

### 2.6 Training data and protocol

#### 2.6.1 SAIR

We train on SAIR, which pairs co-folded protein–ligand structures in ModelCIF format with measured pIC_50_. Structures are read directly from CIF rather than converted to intermediate formats; ligand bond orders are recovered by substructure matching against the deposited SMILES – more reliable than deriving them from a predicted geometry – with fallback to element-only typing when the template cannot be matched. SAIR structures carry no hydrogens, so models trained here are heavy-atom models and are not interchangeable with models trained on protonated data. Preprocessing produced 921,670 usable complexes with no parse failures and no empty records.

Parsed complexes are cached as gzipped shards of atom coordinates and types rather than as materialized graphs. A built graph carries ~600 site atoms of 56-dimensional features and occupies ~195 kB, against 3.3 kB for the parsed form i.e., the difference between 172 GB and 2.9 GB over the full set, and featurization at load time is inexpensive once the structure has been parsed.

#### 2.6.2 Target-disjoint splitting

SAIR contains many ligands per target, a median of 17 and up to 6,750, so a split over complexes places near-identical entries on both sides and the resulting correlation substantially measures memorized targets. **All splits are by protein sequence**. This yields 741,706 training, 89,745 validation and 90,219 test complexes over 3,864, 483 and 483 disjoint targets respectively.

Batches are drawn by a shard-block sampler that shuffles shard order and then shuffles within a block of 16 shards, preserving cache locality; uniform shuffling over 920,000 samples would cost one decompression per sample.

#### 2.6.3 Optimization

AdamW (10^−4^, weight decay 10^−4^); cosine annealing across the epoch budget; batch size 256; dropout 0.3; gradient clipping at norm 1.0; mixed precision; early stopping on validation Pearson *r* with patience 10. Runs terminated at epochs 38–44 of 50 on a single NVIDIA A10G, requiring 39–45 hours. Implementation uses PyTorch 2.0+, PyTorch Geometric, RDKit and BioPython.

### 2.7 Specialized docking modules

i. **Induced-fit docking (PandaDock-Flex)**. Three phases: soft docking with reduced van der Waals radii, receptor side-chain refinement, and full optimization with standard potentials. Receptor strain is tracked separately from the selection energy, so that refinement cannot be rewarded for straining the receptor.
ii. **Metal coordination (PandaDock-Metal)**. Metal centers are identified and coordination geometry assigned (octahedral, tetrahedral, square planar), with distance restraints applied for coordination bonds (M–N 2.0 Å to 2.3 Å, M–O 1.9 Å to 2.2 Å, M–S 2.2 Å to 2.5 Å).
iii. **Tethered docking (PandaDock-Tethered)**. A flat-bottomed harmonic restraint on the ligand centroid,

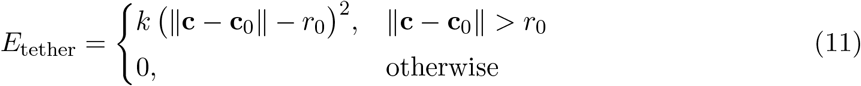

permits free movement within *r*_0_ of the reference and penalizes excursions beyond it.

### 2.8 Evaluation methodology

Because pose accuracy and affinity accuracy fail differently, we evaluate them separately and report controls for each.

1. **Pose evaluation**. Redocking starts from coordinates rebuilt from ligand topology, so crystal geometry cannot leak into the starting pose. RMSD is symmetry-corrected and computed without superposition. We report top-1 accuracy and best-of-*N* accuracy across the returned ensemble; their difference separates selection failure from sampling failure.
2. **Affinity evaluation**. Three controls accompany every correlation. (i) A *variance decomposition* partitioning test label variance into between- and within-target components; the former bounds what a predictor emitting each target’s mean affinity achieves while ignoring the ligand entirely. (ii) A *ligand-only baseline*: ridge regression on sixteen descriptors computed from ligand atoms alone (heavy-atom count, molecular weight, element counts, aromatic and sp^3^ fractions, radius of gyration, maximum extent), trained and evaluated under the identical protocol without ever reading the protein. (iii) *Within-target correlations*, computed per target for targets with at least 20 ligands and then summarized, isolating ligand ranking from target ranking.

Model comparisons are paired by target and tested with a Wilcoxon signed-rank test, since per-target correlations are dispersed enough that comparing summary medians alone can present noise as improvement.

### 2.9 Command-line interface

All functionality is reachable from a unified command-line interface (Table 3) as well as from the Python API.

**Table 3:** Command-line entry points.

| Command | Function |
| --- | --- |
| `pandadock dock` | Flexible-ligand docking (default workflow) |
| `pandadock flex` | Induced-fit docking with receptor flexibility |
| `pandadock metal` | Metal-coordination-aware docking |
| `pandadock tethered` | Restrained docking near a reference pose |
| `pandadock gridbox` | Binding-site box definition |
| `pandadock ml` | Machine-learning-scored workflow |
| `pandadock report` | Interaction analysis and HTML reporting |
| `pandadock-gnn train-sair` | GNN training on the SAIR shard cache |
| `pandadock-gnn predict` | Affinity prediction for a prepared complex |

At inference the binding site is cut to 10 Å around the ligand centroid, matching how the training data was prepared. Passing an entire receptor to a model trained on binding sites produces a confident prediction from an input unlike anything in its training distribution, so the cut is applied by default rather than left to the user.

## 3 Results

### 3.1 Docking benchmark on protein families

Table 4 and Figure 2 report docking on 814 protein–ligand complexes across 14 target families (815 attempted; one RMSD computation failed).

**Table 4:**
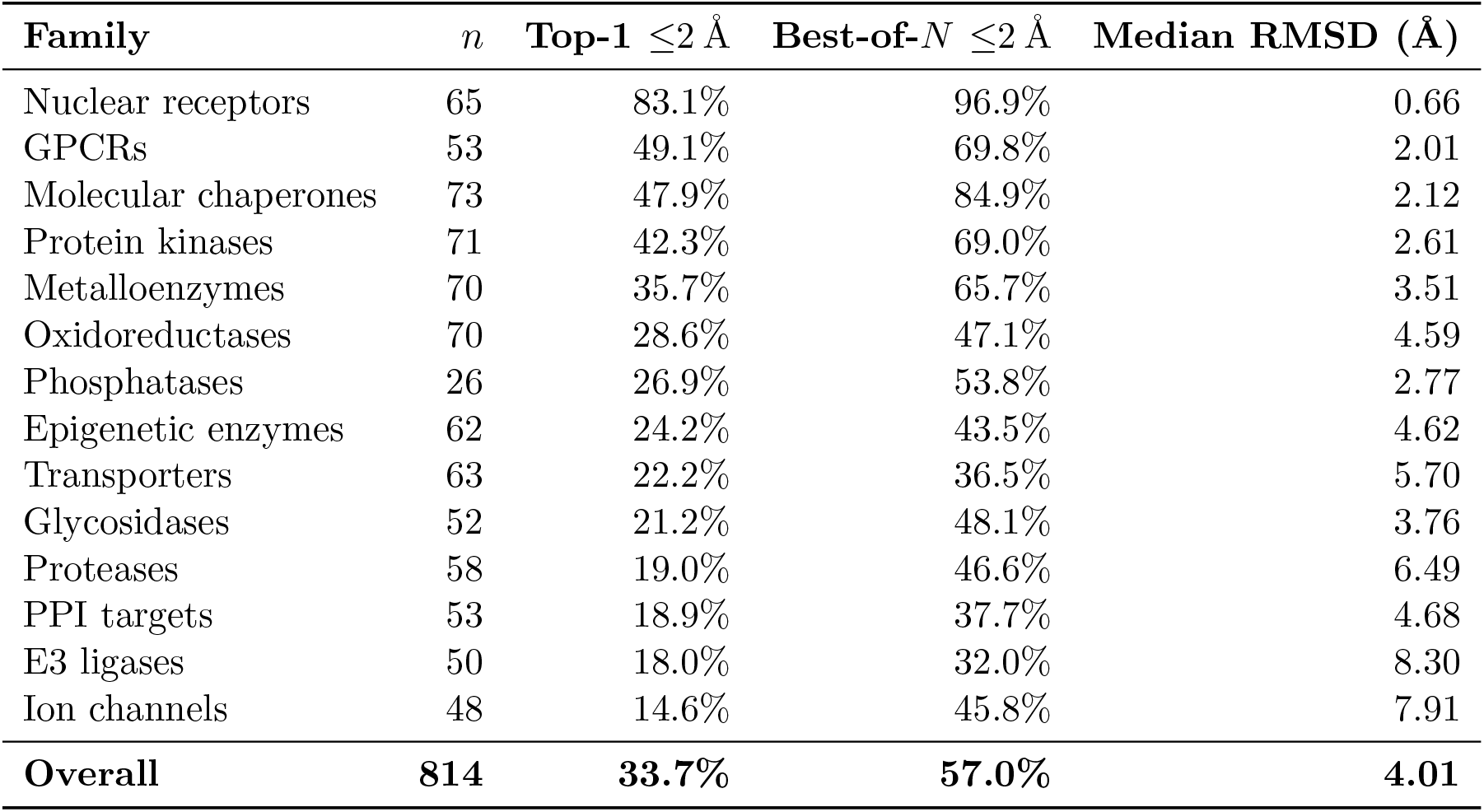
Docking accuracy by target family, 814 complexes. “Top-1” is the highest-ranked pose; “Best-of-*N*” is the closest pose in the returned ensemble.

**Figure 2:**
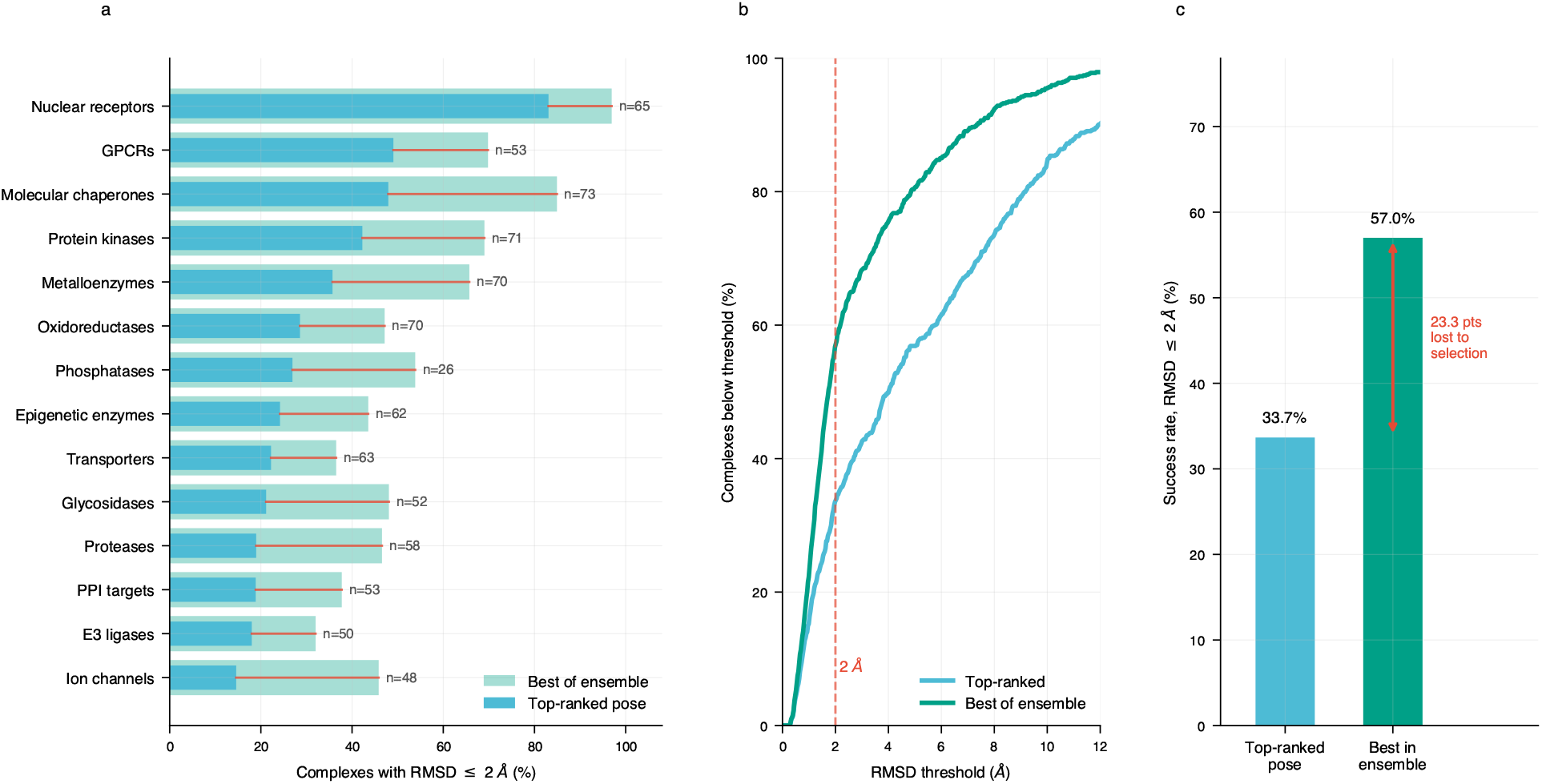
Docking accuracy on 814 complexes. (a) Success rate at 2 Å by target family; bars show the best pose in the returned ensemble (light) and the top-ranked pose (dark), with the connecting line marking the accuracy lost to pose selection. (b) Cumulative accuracy against RMSD threshold. (c) Overall: a correct pose is generated for 57.0% of complexes but ranked first for 33.7%.

Accuracy varies strongly by target class, tracking pocket character. Nuclear receptors, with deep enclosed hydrophobic pockets, reach 83.1% at rank 1, while ion channels and E3 ligases shallow, solvent-exposed and often with protein–protein interface character fall below 20%.

The overall top-1 rate is 33.7%, but a pose within 2 Å is present in the returned ensemble for 57.0% of complexes. The search therefore locates a correct pose substantially more often than the scoring function ranks it first, and the 23.3-point difference is attributable to pose selection rather than to conformational sampling. The gap is non-uniform: molecular chaperones lose 37.0 points between best-of-*N* and top-1 while nuclear receptors lose 13.8. In a separate experiment retaining 20 poses per complex on a 50-complex random sample, the pattern is stronger: 48.0% at rank 1 against an 80.0% ensemble ceiling.

### 3.2 Accuracy and cost against ligand properties

Figure 3 characterizes how both accuracy and runtime respond to ligand size and flexibility. Top-ranked accuracy falls from 46.2% for ligands with a single rotatable bond to 2.4% above twenty, while the ensemble retains a correct pose considerably longer the selection gap persists across the whole range rather than closing at either end. Accuracy peaks for ligands of 20–30 heavy atoms, below which small fragments give few contacts to discriminate on and above which the conformational space grows faster than the sampling budget. Runtime rises with torsion count as intended, since exhaustiveness is set from it, with a median of 334 s and an inter quartile range of 185 s to 506 s.

**Figure 3:**
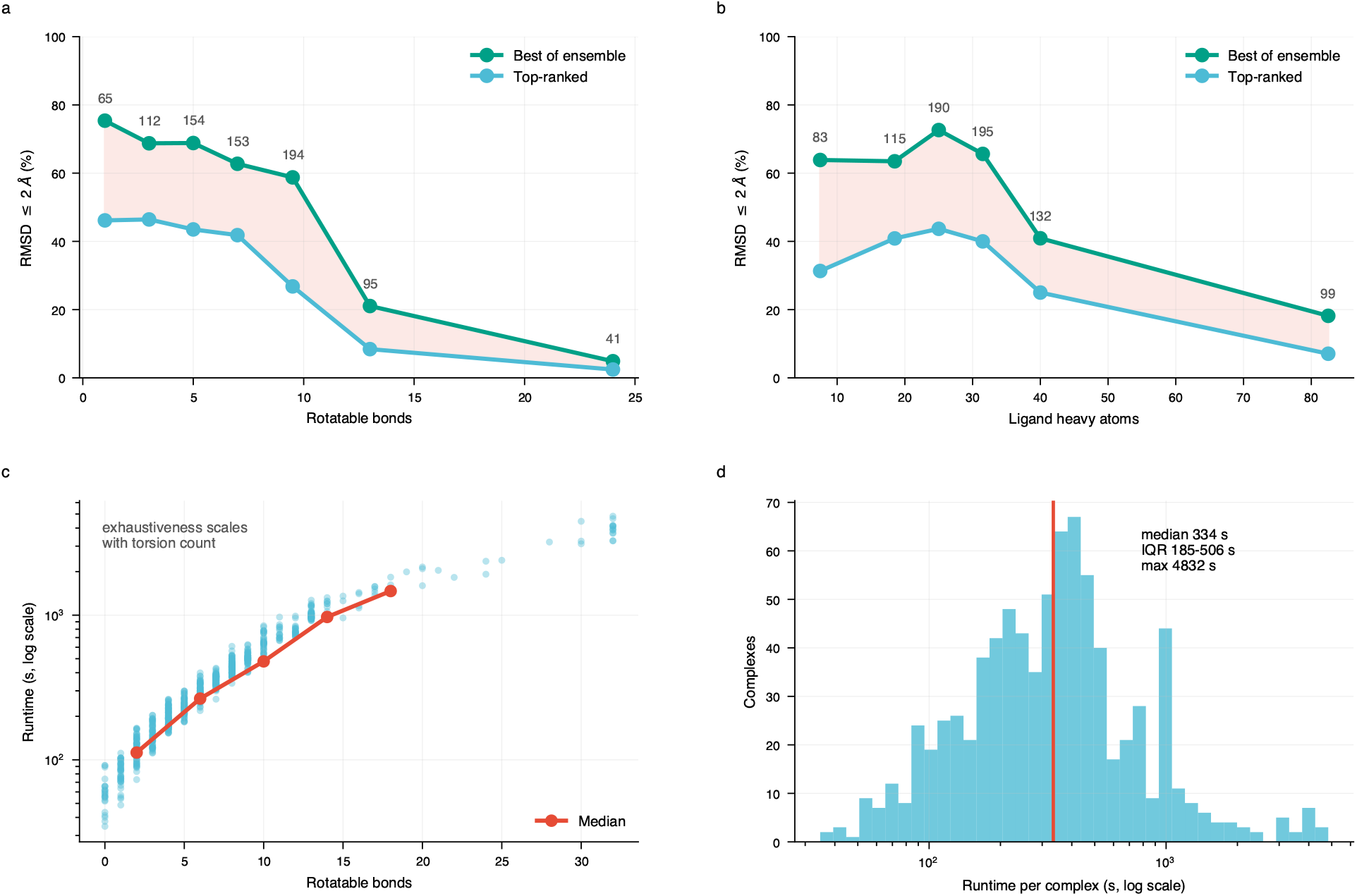
Benchmark characteristics across 814 complexes. (a) Success rate against rotatable-bond count; the shaded band is the accuracy lost to pose selection, and bin populations are annotated. (b) Success rate against ligand heavy-atom count. (c) Runtime against flexibility, with the binned median; exhaustiveness is set from torsion count, so cost rises by design. (d) Distribution of per-complex runtime.

### 3.3 GNN affinity prediction on SAIR

Table 5 and Figure 4 reports the trained model on the 90,219 held-out complexes of the target-disjoint split described in Section 2.6.2, with the controls of Section 2.8. Within-target statistics are computed over the 234 targets carrying at least 20 ligands, comprising 88,880 complexes – 98.5% of the split, so the statistic is representative rather than a favorable subset.

**Table 5:** Affinity prediction on the held-out SAIR test split, with controls.

| Predictor | Pooled $r$ | Median within-target $r$ |
| --- | --- | --- |
| Target-mean predictor (ignores ligand) | 0.541 | — |
| Ligand descriptors only (ignores protein) | 0.383 | 0.196 |
| **PandaDock-GNN** | **0.407** | **0.237** |

**Figure 4:**
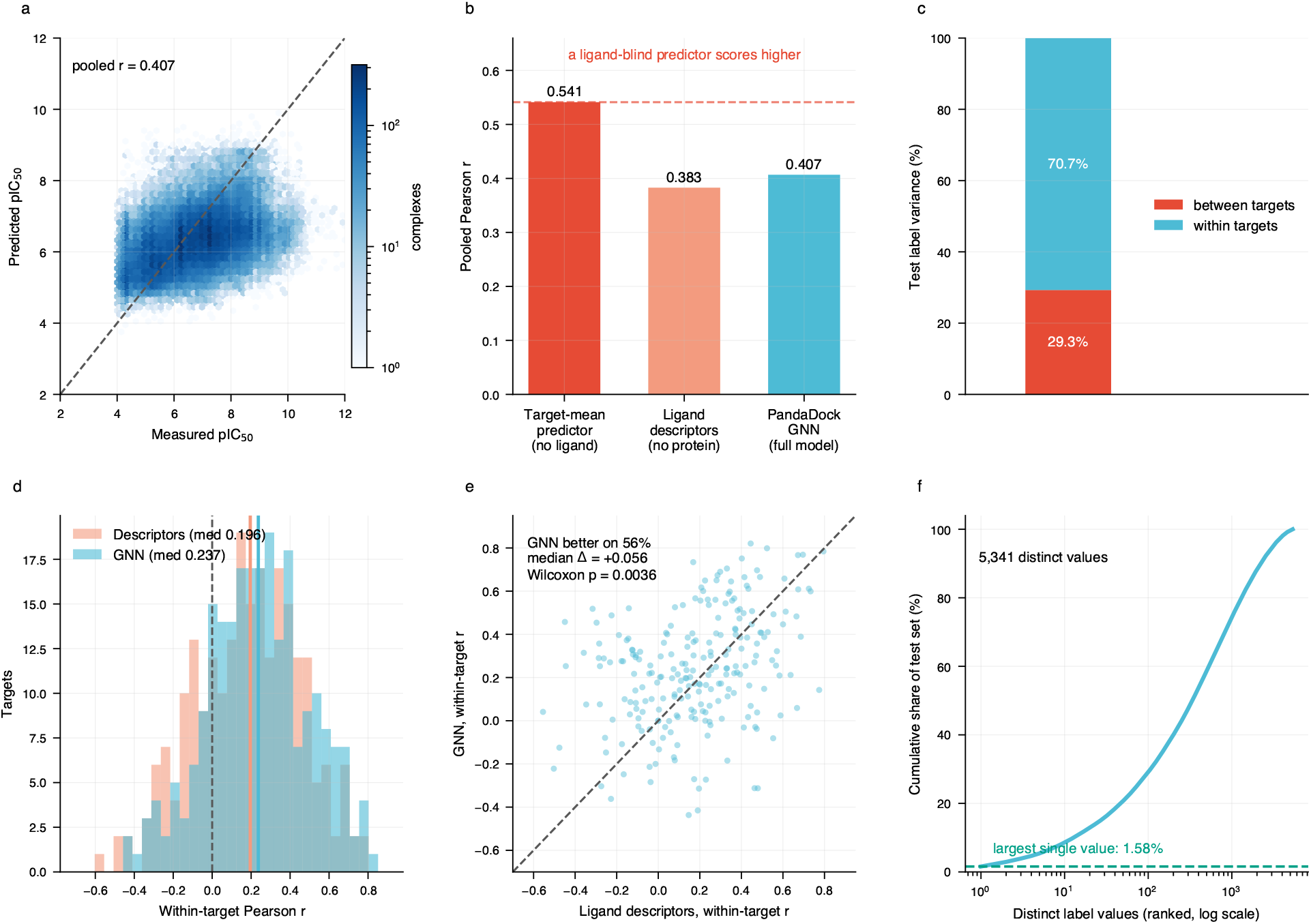
Affinity prediction on held-out SAIR with evaluation controls. (a) Predicted against measured pIC_50_ for 90,219 complexes over 483 disjoint targets. (b) Pooled correlation against two controls: a target-mean predictor that never sees the ligand scores higher than the trained network. (c) 29.3% of test label variance lies between targets, which is what makes the target-mean control competitive. (d) Within-target correlations for the network and for ligand descriptors alone, over the 234 targets with at least 20 ligands. (e) The same comparison paired by target. (f) Test labels are well distributed, so these correlations are not artefacts of censoring (contrast Figure 10).

Test labels are well distributed; 5,341 distinct values with no value exceeding 1.58% of the split, so these correlations are not artifacts of a collapsed label distribution.

Two observations follow from the controls. First, 29.29% of test label variance lies between targets, so a predictor emitting each target’s mean while ignoring the ligand entirely would score a pooled *r* of 0.541, above the model’s 0.407. Pooled correlation on a multi-target split cannot by itself demonstrate ligand discrimination, and the within-target figure is the appropriate headline. Second, ligand descriptors alone reach a median within-target *r* of 0.196 against the network’s 0.237; paired across 234 targets the difference is reliable (Wilcoxon *p* = 0.0036; better on 56.0% of targets; median +0.056), though the bootstrap interval on the median change spans zero ([−0.002, +0.098]), so the direction is established while the magnitude is not.

### 3.4 Transfer to experimental crystal structures

The model is trained entirely on co-folded predicted structures. To test whether this transfers to experimental geometry, we evaluated it on 202 crystal structures spanning 13 families with measured *K*_*i*_ (*n* = 53), IC_50_ (*n* = 117), *K*_*d*_ (*n* = 12) or EC_50_ (*n* = 20), using the same site extraction and hydrogen handling as at inference (Table 6 and Figure 5).

**Table 6:** Affinity correlation on 202 experimental crystal structures with measured binding constants.

| Family | $n$ | $r$ | Family | $n$ | $r$ |
| --- | --- | --- | --- | --- | --- |
| Glycosidases | 5 | 0.815 | Oxidoreductases | 18 | 0.204 |
| Transporters | 23 | 0.586 | Ion channels | 13 | 0.199 |
| Nuclear receptors | 22 | 0.365 | Chaperones | 20 | 0.085 |
| Metalloenzymes | 24 | 0.351 | GPCRs | 31 | 0.022 |
| Kinases | 19 | 0.335 | Epigenetic | 5 | 0.002 |
| Proteases | 14 | 0.305 | E3 ligases | 6 | -0.157 |
| **Overall:** $r = 0.467$, $\rho = 0.475$, $n = 202$ | | | | | |

**Figure 5:**
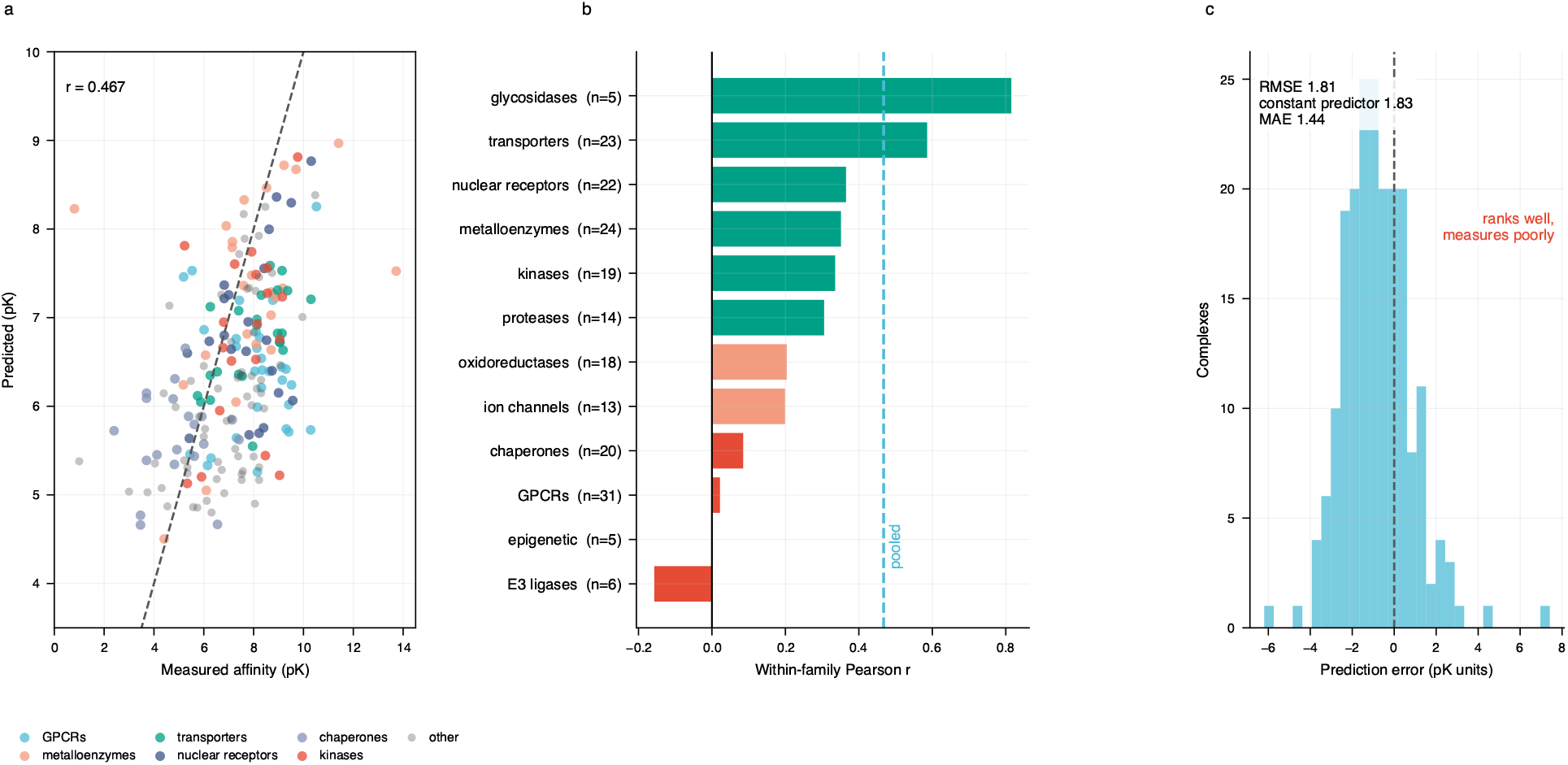
Transfer to experimental crystal structures. (a) Predicted against measured affinity for 202 crystal complexes with reported *K*_*i*_, *K*_*d*_, IC_50_ or EC_50_, coloured by the six largest families. Within-family correlations, sorted; the dashed line marks the pooled value, which averages over markedly heterogeneous behavior. (c) Error distribution: RMSE is barely below that of a constant predictor, so the model ranks considerably better than it measures.

Pooled correlation is *r* = 0.467, higher than on the model’s own test split, indicating that transfer from co-folded to experimental geometry holds. Two qualifications apply. Per-family correlations span −0.157 to 0.586, with the largest family (GPCRs, *n* = 31) at 0.022, so the pooled value averages over heterogeneous behavior and should be read alongside the breakdown. And absolute accuracy is limited: RMSE is 1.811 against 1.827 for a constant predictor, with MAE 1.438 log units, and predictions span 4.50–8.97 where measurements span 0.81–13.72. The model ranks considerably better than it measures; its output is compressed toward the training mean and requires calibration before absolute values are reported.

### 3.5 A prospective within-target series: GABA_*A*_ receptor ligands

The crystal-structure and SAIR test-split results above are pooled across many targets, and Section 2.8 showed that pooled correlation on such a set is not evidence of ligand ranking. A direct test of ligand ranking needs a series of compounds against one fixed target with poses generated independently of any of the models being compared.

We assembled such a series: 30 GABA_*A*_ receptor ligands (a propofol/etomidate chemotype series plus reference general anesthetics) with measured EC_50_ against a single receptor structure, none of which appear in SAIR, BindingDB or PDBbind. Ligands were supplied as unposed three-dimensional conformers and docked into the receptor with the default algorithm which is protein-ligand flexible docking (Algorithm 1); the top-ranked pose from that single docking run was then scored by every model under comparison, so all methods are compared on identical geometry. Twenty independently obtained docking-based scores for the same series, spanning three scoring functions (Vina, Vinardo, AutoDock4) under rigid and flexible protocols on CPU and GPU, provide external context for where PandaDock’s own scores fall.

PandaDock’s empirical scoring function reaches *r* = 0.768 (*ρ* = 0.656), ranking 8th of 25 methods evaluated on this series: above all four AutoDock Vina configurations and all four Vinardo configurations, below only the AutoDock4-GPU variants (Table 7). This is an independent, single-target confirmation of the redocking result – the search and empirical scoring components are competitive with established tools.

**Table 7:** Selected methods on the 30-compound GABA_*A*_ series, ranked by Pearson *r* against pEC_50_. AutoDock4-GPU rows are independently obtained runs under varying rigid/flexible and interaction-term settings, included for external context; the full 25-method ranking is Figure 6d.

| Method | Pearson $r$ | Spearman $\rho$ |
| --- | --- | --- |
| AutoDock4-GPU (rigid, intermolecular) (external) | 0.857 | 0.827 |
| AutoDock4-GPU (rigid) (external) | 0.822 | 0.827 |
| **PandaDock (empirical)** | **0.768** | **0.656** |
| Vinardo (rigid, intermolecular) (external) | 0.752 | 0.528 |
| AutoDock Vina (rigid) (external) | 0.673 | 0.489 |
| GNN, ULVSH+BindingDB+PDBBind | 0.481 | 0.393 |
| GNN, BindingDB | 0.364 | 0.148 |
| GNN, SAIR (this paper’s model) | 0.356 | 0.406 |
| GNN, BindingDB+ULVSH | 0.351 | 0.528 |
| AutoDock4 (rigid) (external) | 0.348 | 0.008 |

Every GNN variant we evaluated, including the SAIR model reported as this paper’s primary result, scores below all four AutoDock Vina configurations on this series: *r* = 0.356 for SAIR, against 0.673 for AutoDock Vina (rigid). This is consistent with, and independent confirmation of, the within-target ceiling identified on SAIR’s own held-out complexes (median *r* = 0.237, Table 5) and on the crystal set’s largest family (GPCRs, *r* = 0.022, Table 6): four separate measurements, on four disjoint sets of targets and compounds, place the GNN’s ligand-ranking ability at or below what classical empirical scoring already achieves. We are not aware of a within-target series on which any GNN variant evaluated here outperforms Vina.

The output-range pattern of Figure 4 reproduces exactly: the model trained on BindingDB alone spans 2.25 pK units across these 30 predictions, falling to 0.58 once ULVSH is added to training and partially recovering to 1.21 once PDBbind is added as well (Figure 6e). ULVSH’s censored labels compress a model’s output regardless of what else it is trained on, and the effect is visible on a held-out series the model has not seen in any form.

**Figure 6:**
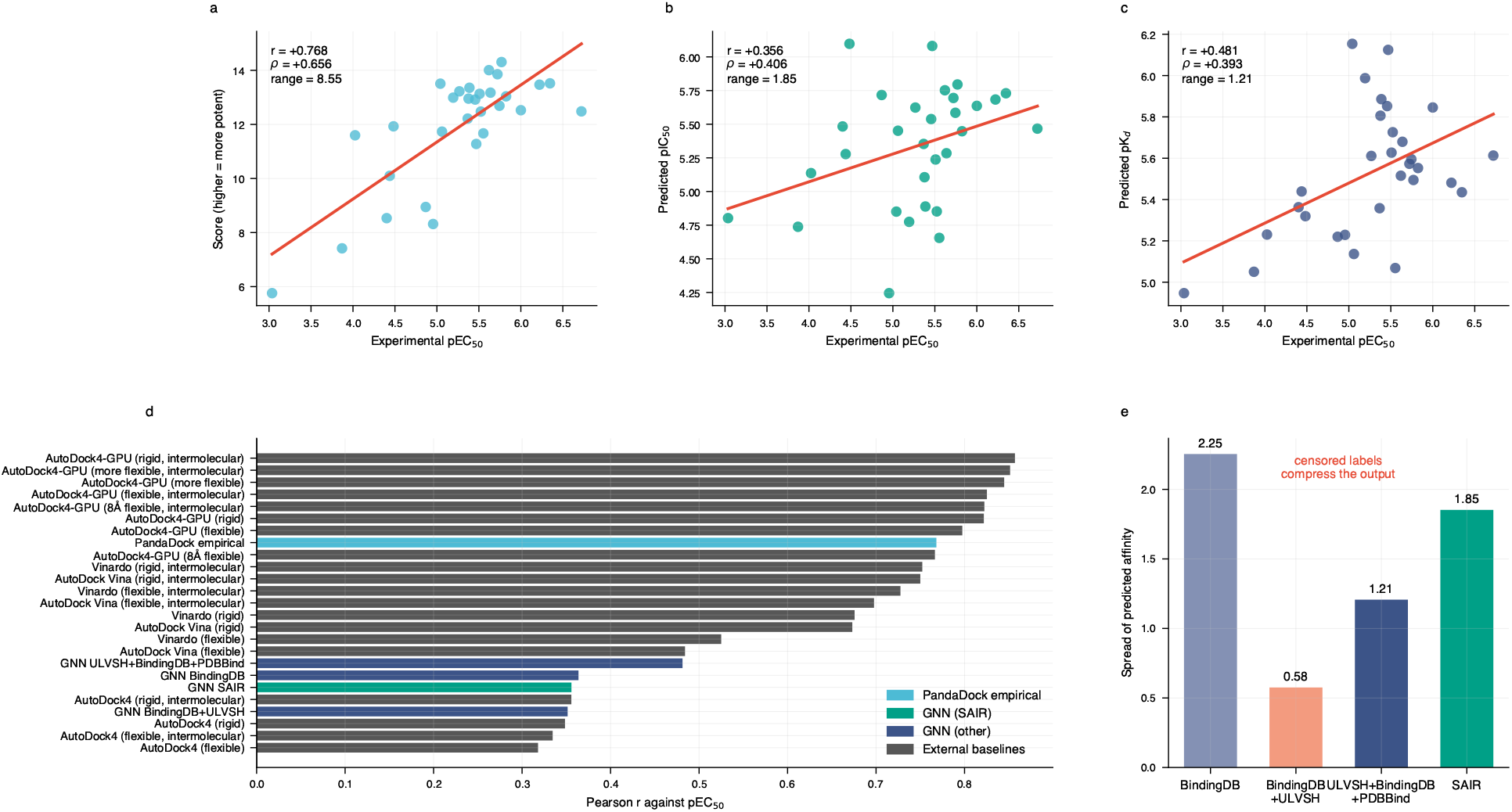
A 30-compound GABA_*A*_ receptor series, one target, identical poses for every method. (a) PandaDock’s empirical scoring function. (b) The GNN trained on SAIR – the model reported elsewhere in this paper. (c) The GNN trained on BindingDB, ULVSH and PDBbind combined. (d) Pearson *r* for every method on the same 30 compounds; PandaDock’s empirical score is competitive with established docking scoring functions, while every GNN variant, SAIR included, scores below every Vina and Vinardo configuration tested. (e) Output dynamic range by training data: the ULVSH-containing models are compressed, consistent with Figure 10.

### 3.6 Validation at scale on the PDBbind v2020 refined set

Beyond the crystal-structure and GABA_*A*_ checks above, we evaluated every model against the PDBbind v2020 refined set (5,316 complexes with curated − log(*K*_*d*_*/K*_*i*_) labels), which is roughly 26*×* larger than the crystal-structure check reported elsewhere in this paper and lets us test generalization at a scale those smaller checks cannot.

No docking was required: PDBbind provides each complex’s co-crystallized ligand pose, so this evaluates affinity prediction on native geometry, using the same 10 Å site extraction and hydrogen handling as every other affinity result in this paper. 4,640 of 5,316 complexes parsed successfully; the remainder failed RDKit sanitization on a minority of ligands with unusual valences, a known limitation of that parser rather than a property of any particular subset of the data. Labels are not degenerate: 816 distinct values across the parsed set, with no single value exceeding 0.9%, so this dataset does not exhibit the censoring failure identified in Section 4.2.

One methodological asymmetry matters throughout this section: **ULVSH+BindingDB+ PDBBind was trained on this dataset**, so its result here is not a held-out evaluation and should be read as an upper bound inflated by an unknown degree of train/test overlap. BindingDB, BindingDB+ULVSH, and the SAIR model reported throughout the rest of this paper never saw PDBbind in any form; for these three the result is a fully independent, large-scale evaluation additional to the crystal-structure and GABA_*A*_ checks above.

Figure 7 and Table 8 report pooled and within-target correlation for all five models. The independent SAIR model reaches *r* = 0.531 (*ρ* = 0.523), higher than ULVSH+BindingDB+PDBBind’s *r* = 0.387 despite it having trained on this exact data – genuinely held-out performance on this dataset is therefore worse than 0.387, not better, for any model that mixes PDBbind with other training sources.

**Table 8:** PDBbind v2020 refined set. Within-target *r* uses the 103 receptor-sequence targets with ≥ 5 complexes. All rows except the last are scored on the full pooled set (n = 4,640), which includes complexes ULVSH+BindingDB+PDBBind and PDBbind-only were trained or validated on. The PDBbind-only (held-out) row instead uses only that model’s own disjoint test split (n = 624, never seen during its training) and is the only row in this table free of train/test overlap with PDBbind.

| Model | Pooled $r$ | Pooled $\rho$ | Median within-target $r$ |
| --- | --- | --- | --- |
| SAIR (fully independent) | 0.531 | 0.523 | 0.305 |
| ULVSH+BindingDB+PDBBind (trained) | 0.387 | 0.382 | 0.318 |
| PDBbind-only (trained on this data) | 0.713 | 0.721 | 0.621 |
| BindingDB (independent) | 0.277 | 0.281 | 0.240 |
| BindingDB+ULVSH (independent) | -0.079 | -0.022 | 0.126 |
| **PDBbind-only (held-out, $n = 624$)** | **0.690** | **0.674** | – |

**Figure 7:**
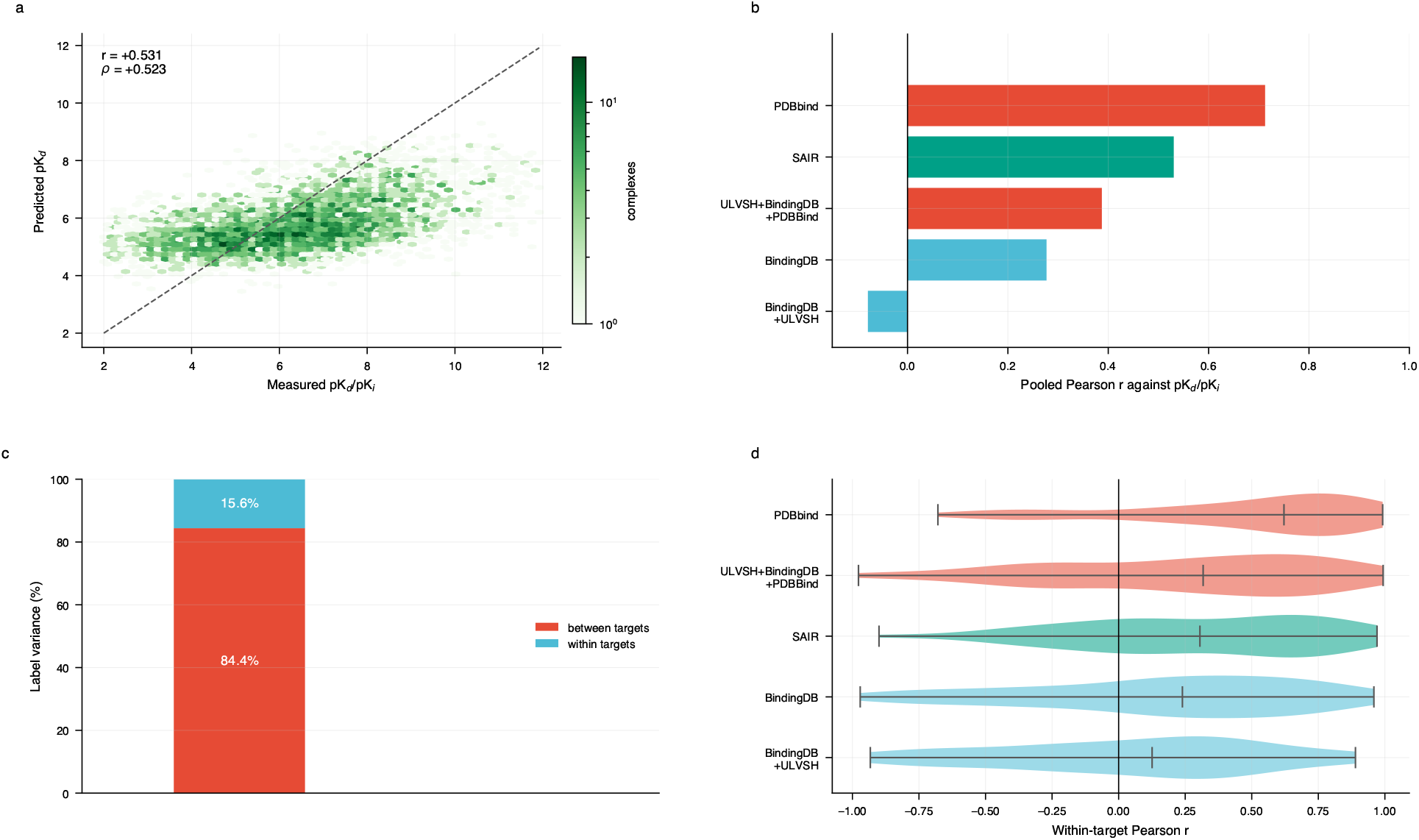
PDBbind v2020 refined set, native crystal poses, n = 4,640. (a) The SAIR model – fully independent of this dataset – against measured affinity. (b) Pooled Pearson *r* for all five models; red bars trained on some or all of this data, green fully independent, blue independent of PDBbind specifically. (c) 84.4% of label variance lies between the 2,963 distinct receptor-sequence targets, higher than on SAIR’s own test split, consistent with PDBbind’s structure of many single-complex targets. (d) Within-target Pearson *r* for targets with at least 5 complexes (103 targets, 1,134 complexes); a 2-complex group is excluded because its correlation is *±*1 by construction and not a statistic.

We additionally trained a GNN on PDBbind alone, under a target-disjoint split (3,553 complexes over 2,370 training targets; 463/296 validation; 624/297 test, split by exact receptor sequence so no target’s complexes appear in more than one split). Scored on the same pooled n = 4,640 set as the other four models – 4,016 of which it has seen in training or validation – it reaches *r* = 0.713, the highest pooled figure of the five, but this number is inflated by that overlap in the same way ULVSH+BindingDB+PDBBind’s is. The number that reflects genuine generalization is its performance on its own 624 held-out test complexes, which the model never saw in any form during training: there it reaches *r* = 0.690 (*ρ* = 0.674, RMSE = 1.51 pK units) – still higher than every independent model in the table, and consistent with a properly disjoint-split model trained specifically on this data reaching a materially higher correlation than a model trained on other sources and merely applied to it.

Two further observations. First, SAIR’s within-target median of 0.305 on 4,640 complexes is consistent with its own held-out test split (0.237, Table 5) and with the GABA_*A*_ series (0.356, Table 7) – three measurements on three disjoint sets of targets and compounds now agree to within 0.07 of each other, which is the strongest evidence in this paper that the within-target ceiling is a property of the model rather than of any one evaluation set. Second, BindingDB+ULVSH’s negative pooled *r* (−0.079) despite a positive within-target median (0.126) is the output-compression artefact of Figure 4 operating at a scale where it visibly breaks the pooled statistic rather than merely narrowing it, a more severe expression of the same failure documented on the GABA series and the SAIR test split.

Taken together, these results show that PandaDock’s affinity predictions generalize to structures and targets never seen in training, at full scale and with the training/held-out distinction made explicit throughout: a fully independent model (SAIR) reaches *r* = 0.531 on 4,640 unseen complexes, and a model trained specifically on PDBbind reaches *r* = 0.690 on its own target-disjoint held-out test set.

### 3.7 Suitability for pose rescoring

Because the platform offers a hybrid workflow in which learned scoring may rerank generated poses, we evaluated whether the affinity model improves pose selection. For 50 complexes we generated 20 poses each and compared three selections: the pose ranked first by the empirical function, the pose ranked first by the GNN, and the lowest-RMSD pose available (Table 9 and Figure 8).

**Table 9:** Pose selection on 50 complexes, 20 poses each. “Ensemble best” bounds what any rescorer could achieve on this pose set.

| Selection | Median RMSD (Å) | $\leq 2$ Å | $\leq 1$ Å |
| --- | --- | --- | --- |
| Empirical scoring function | 2.09 | 48.0% | 24.0% |
| GNN rescoring | 5.36 | 22.0% | 14.0% |
| Ensemble best | 1.26 | 80.0% | 38.0% |

**Figure 8:**
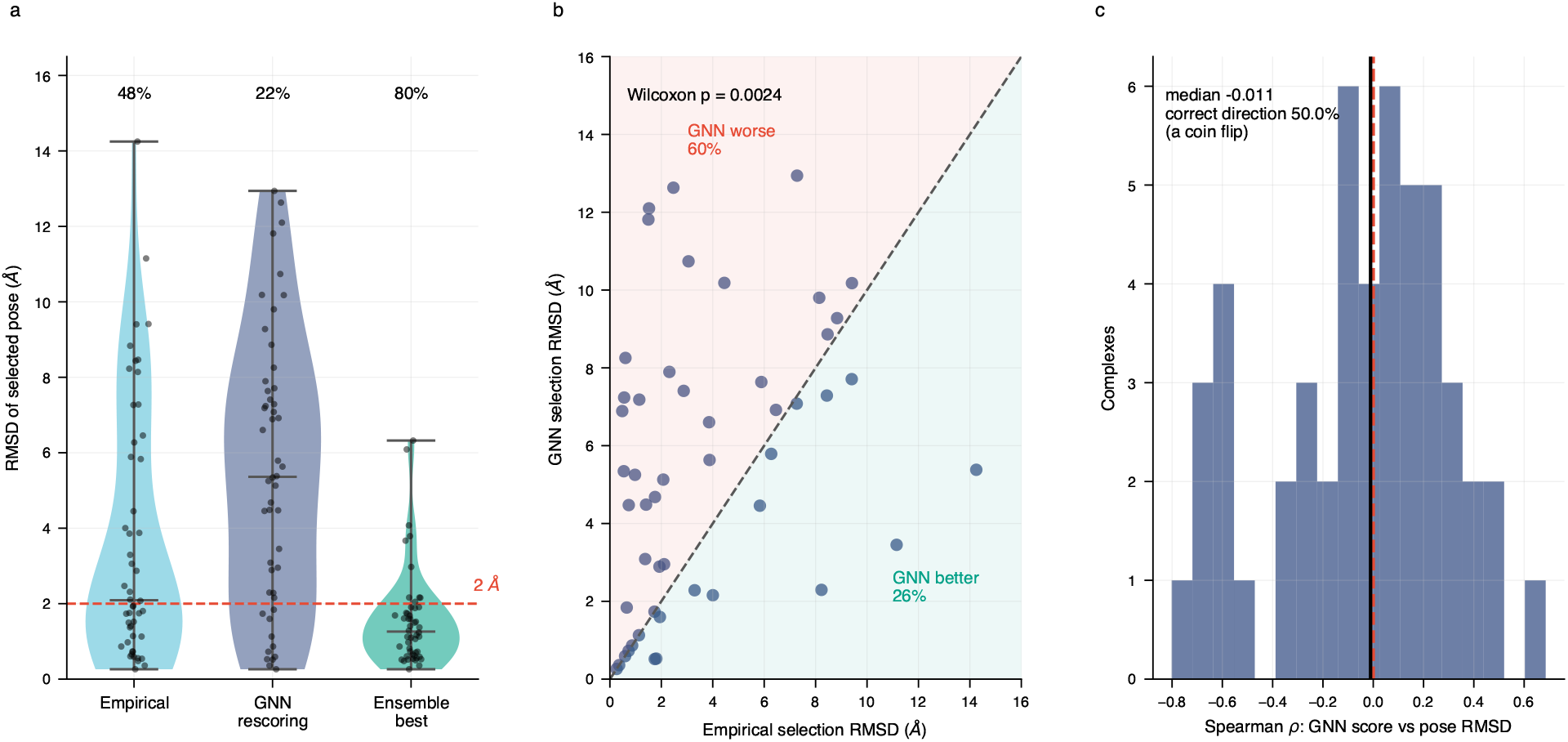
The affinity model does not rank poses. (a) RMSD of the pose selected by each method across 50 complexes, with the percentage within 2 Å above each distribution. (b) Paired by complex: points above the diagonal are complexes where GNN rescoring chose a worse pose. (c) Per-complex rank correlation between GNN score and pose RMSD; a working rescorer would produce a distribution centered well below zero.

GNN rescoring is reliably worse: it selects a poorer pose on 60.0% of complexes and a better one on 26.0%, with a median RMSD change of +0.905 Å (Wilcoxon *p* = 0.0024). The per-complex rank correlation between GNN score and pose RMSD has median −0.011 and points in the correct direction on 50.0% of complexes.

The model is trained exclusively on near-native co-folded poses and has never encountered incorrect geometry, so it has no representation with which to penalize it and assigns out-of-distribution poses arbitrary rather than poor scores. Pose discrimination requires training against decoys at known RMSD, which is a different learning problem from affinity regression. **We therefore recommend the empirical function for pose ranking and the GNN for affinity estimation on an already-selected pose**, and the software defaults accordingly.

A simpler test is misleading here and we note it as guidance. Rigidly displacing a crystal ligand and rescoring suggests strong pose sensitivity; the model marks down 71.3% of 1 Å and 73.3% of 5 Å displacements, but translation reduces contact count in an easily detected way, whereas docked decoys are physically plausible poses preserving contacts. Displacement tests should not be taken as evidence of pose discrimination.

### 3.8 Objective ablation

Since within-target ranking is the ability of interest, and regression on absolute affinity is dominated by the between-target component, we added a loss term removing each target’s batch mean from both prediction and label, retaining the absolute loss so that the model still emits an absolute affinity. Batch composition supported it: 94% of a typical batch belonged to a target with at least two complexes present.

Median within-target *r* moved from 0.208 to 0.237. Paired across 234 targets this is not distinguishable from noise (Wilcoxon *p* = 0.4968; better on 54.3% of targets; bootstrap interval on the median change [−0.013, +0.036]). We report the ablation as negative. The inter quartile range of per-target correlations is approximately 0.35, so a median shift of 0.03 lies well inside the spread, and the intervention would have appeared as a 14% relative improvement had summary medians been compared without a paired test.

### 3.9 Computational performance

Median redocking runtime was 334 s per complex at default settings on CPU. Table 10 and Figure 9 summarizes the effect of Algorithm 2.

**Table 10:** Affinity grid construction, dense versus blocked evaluation. Grids are bit-identical in every case (maximum absolute difference 0.0).

| Complex | Dense (s) | Blocked (s) | Speedup |
| --- | --- | --- | --- |
| 2wi3 | 205.0 | 29.4 | 7.0× |
| 2jjc | 103.6 | 18.6 | 5.6× |
| 5fpe | 67.7 | 7.0 | 9.7× |
| 2wi2 | 104.2 | 15.4 | 6.8× |
| **Total** | **480.5** | **70.4** | **6.8×** |

**Figure 9:**
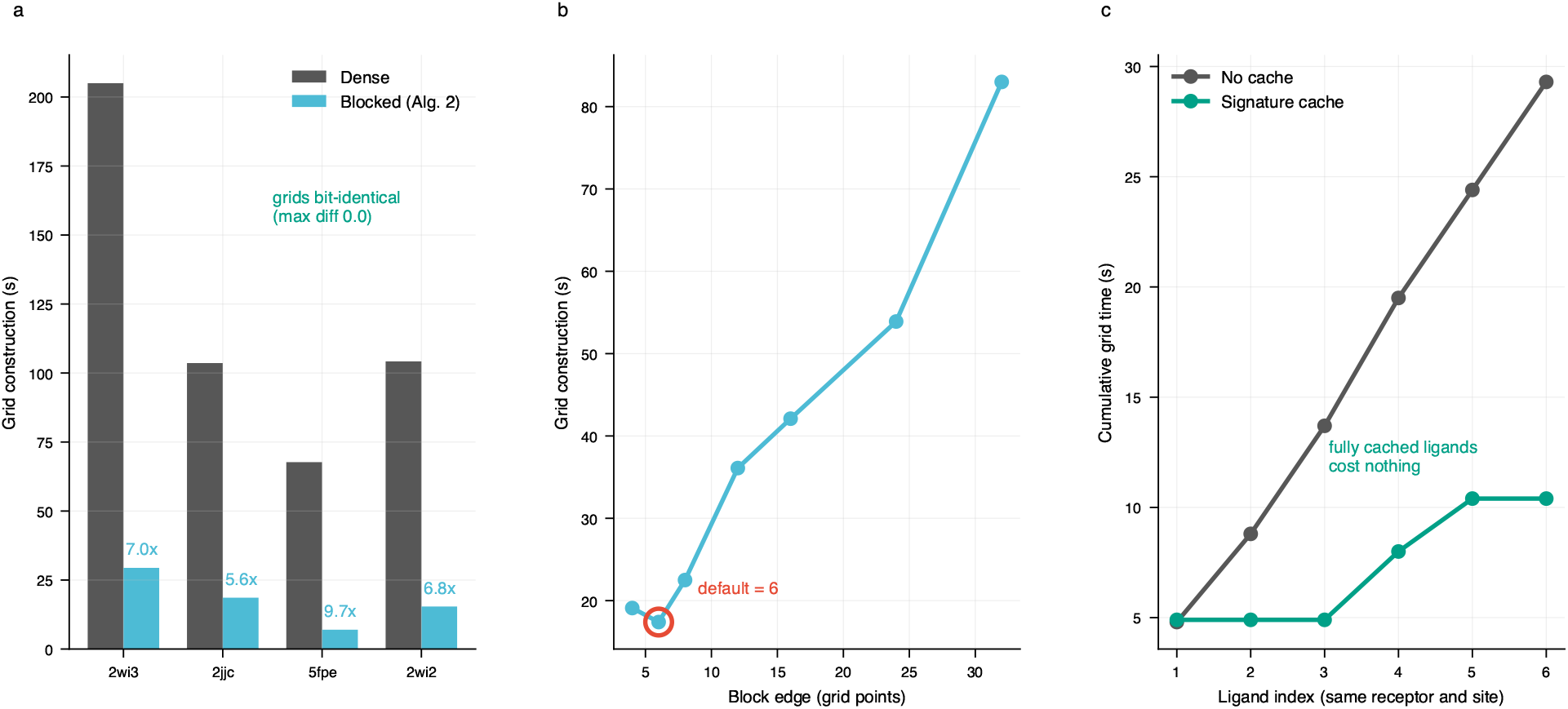
Affinity grid engine. (a) Blocked neighbor selection against dense evaluation on four receptors; grids are bit-identical in every case. (b) Block size against construction time, measured on a 65^3^ grid and a 5,800-atom receptor; the default is the measured minimum. (c) Cumulative grid cost across six ligands docked into one receptor, with and without the signature cache.

**Figure 10:**
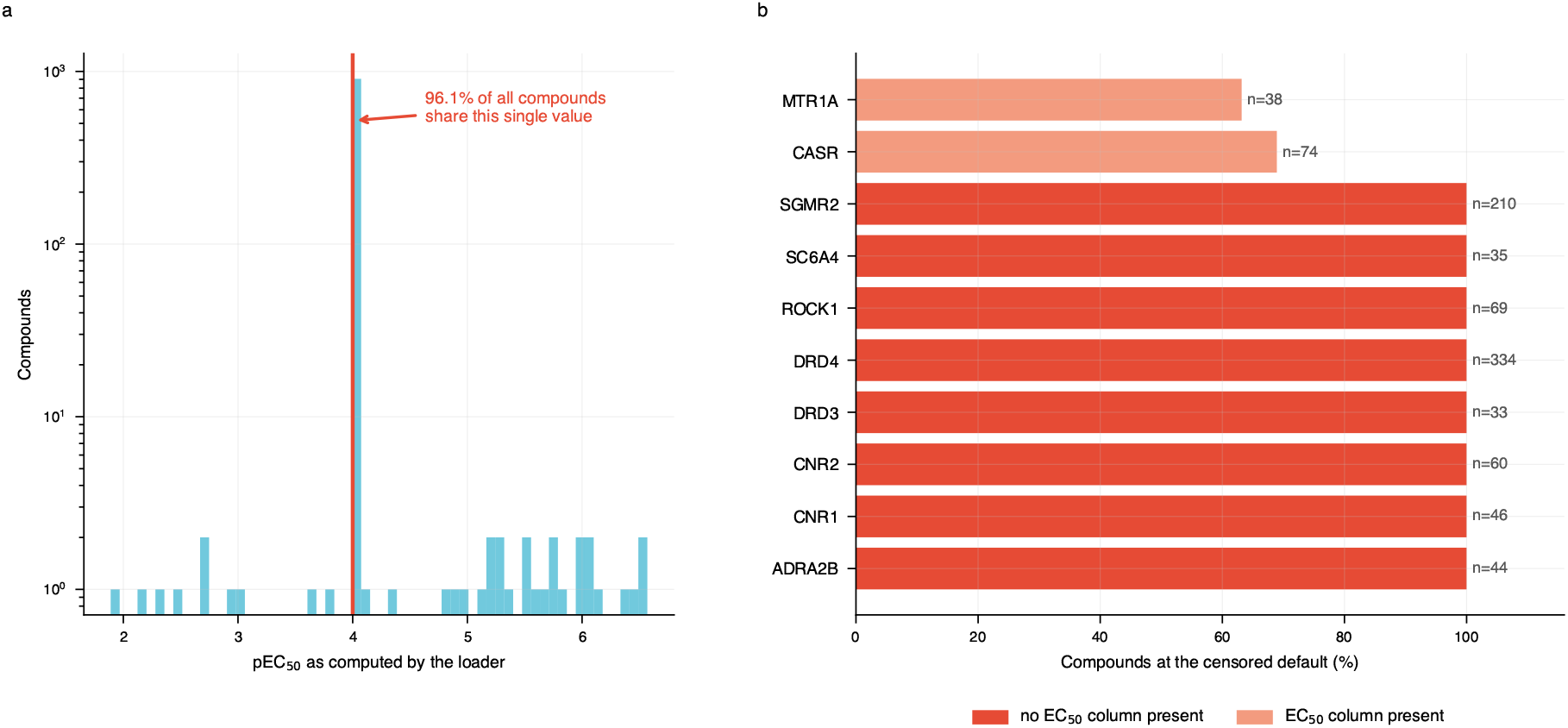
A censored-label artefact. (a) Label distribution of the withdrawn benchmark as computed by its loader; 96.1% of compounds share the single censored value at pEC_50_ = 4.0 (note the logarithmic count axis). (b) Share of compounds at that default per target: eight of ten targets expose no column the loader recognises as EC_50_ and are therefore entirely censored.

End to end this reduces a representative docking run from approximately 274 s to 99 s. Grid caching across a six-ligand series against one receptor reduces grid construction from 29.3 s to 10.4 s, with fully cached ligands incurring no grid cost; the benefit grows with campaign size, since the cache converts a per-ligand cost into a one-time cost.

The remaining cost is the search itself: 168,754 energy-and-gradient evaluations for a single complex at exhaustiveness 8, each over approximately 50 atoms, dominated by interpreter overhead rather than arithmetic. Batching the independent Monte Carlo runs into single tensor operations the strategy underlying GPU-accelerated dockers [Santos-Martins et al., 2021; Tang et al., 2022] is the natural next step and is not implemented in the current version.

## 4 Discussion

### 4.1 Where accuracy is lost

The redocking results locate the dominant error precisely. A pose within 2 Å is generated for 57.0% of complexes but ranked first for 33.7%, and on the 20-pose subset the gap widens to 80.0% against 48.0%. Conformational sampling is therefore not the binding constraint at these settings; pose ranking is. This has a practical consequence for users, who should inspect the returned ensemble rather than the top pose alone, and a development consequence: effort directed at ranking has substantially more headroom than effort directed at sampling.

Our attempt to close that gap with the learned scoring function failed, and informatively. An affinity model is not automatically a rescoring function: the two tasks differ in what they must represent – one the relationship between a native complex and its binding constant, the difference between correct and incorrect geometry, and a model trained only on near-native structures has no exposure to the latter.

### 4.2 Interpreting learned scoring functions

The controls of Section 2.8 are inexpensive and we recommend them generally. On our test split a predictor ignoring the ligand entirely achieves a higher pooled correlation than the trained network, which means a pooled figure reported alone would be uninterpretable. Ligand descriptors reach 83% of the network’s within-target performance, bounding the structural contribution at approximately 0.04 in correlation – real, but far smaller than a pooled number suggests.

We also encountered a labelling artefact worth recording. During development we evaluated on a public virtual-screening dataset and obtained a Pearson *r* of 0.82, apparently far above published baselines. On inspection, that dataset stores assay results in target-specific columns (*K*_*i*_ in nM, percentage inhibition, percentage displacement); only two of ten targets expose a column parseable as EC_50_, and unparseable values fall back to a censored default. The consequence is that 96.1% of compounds share a single label and only 37 of 943 carry a distinct measured value. We withdraw that result. The failure is silent — no error is raised and training curves appear healthy, and we recommend that affinity benchmarks routinely report the number of distinct label values and the share held by the most common.

A related integrity practice applies to PDBbind evaluation more broadly. Benchmarking scripts for affinity correlation are easy to get subtly wrong in ways that inflate reported numbers without any error being raised: silently sub sampling to a small, non-random subset of a dataset while reporting the full size; applying a post-hoc scale correction that selects, per algorithm, whichever of several transformations – including a linear rescaling fitted to the exact evaluation labels – maximizes agreement with those same labels before any correlation is computed; or including a mock-evaluation mode in which the “prediction” is generated as the true label plus Gaussian noise, left enabled by default. Any one of these invalidates a correlation computed from it. We report the checks in Section 3.6 at full dataset scale (n = 4,640), with no post-hoc rescaling and no mock-evaluation path, and recommend the same practice – full-scale evaluation, a fixed prediction pipeline, and no label-informed transform – as a general safeguard for learned scoring functions.

### 4.3 Limitations

Redocking uses a box centred on the crystal ligand, which is standard but optimistic relative to blind docking. The GNN is trained on co-folded predicted structures whose within-target pose diversity may not reflect experimental reality, and we have not separated this from model capacity as an explanation for the modest within-target ceiling. The pose-rescoring experiment cuts the binding site once around the crystal ligand, so poses displaced far from it are scored on an unrealistic input; the conclusion is unchanged on complexes whose poses lie close together, but the magnitude may be overstated. The crystal transfer set is modest at 202 complexes, with several families below *n* = 10. Absolute affinity predictions are uncalibrated. The search is CPU-only in the current version of the release. GPU based docking is planned for future releases.

## 5 Conclusion

PandaDock provides a complete open-source docking platform: flexible-ligand search with analytic SO(3) gradients, an exact blocked grid engine with screening-oriented caching, specialized induced-fit, metal and tethered modules, and an SE(3)-equivariant scoring function trained on 741,706 complexes. On 814 complexes across 14 families it generates a sub-2 Å pose for 57.0% and ranks one first for 33.7%; the learned scoring function transfers from predicted to experimental structures at *r* = 0.467 across 202 crystal complexes with measured binding constants. We report the evaluation controls that make those numbers interpretable and document where the platform does not perform, notably that the affinity model should not be used to rescore poses. The software, the benchmarking harness and the per-complex results underlying every figure quoted here are openly available.

## Data and Code Availability

PandaDock is available at https://github.com/pritampanda15/PandaDock under an open-source license and installable from PyPI. The benchmarking harness — redocking, label auditing, variance decomposition, ligand-only baseline, paired model comparison and pose rescoring — is in `benchmarking/`, with per-complex predictions underlying every correlation reported here.

